# Unraveling the mechanism of the cadherin-catenin-actin catch bond

**DOI:** 10.1101/306761

**Authors:** Shishir Adhikari, Jacob Moran, Christopher Weddle, Michael Hinczewski

## Abstract

The adherens junctions between epithelial cells involve a protein complex formed by E-cadherin, *β*-catenin, *α*-catenin and F-actin. The stability of this complex was a puzzle for many years, since in vitro studies could reconstitute various stable subsets of the individual proteins, but never the entirety. The missing ingredient turned out to be mechanical tension: a recent experiment that applied physiological forces to the complex with an optical tweezer dramatically increased its lifetime, a phenomenon known as catch bonding. However, in the absence of a crystal structure for the full complex, the microscopic details of the catch bond mechanism remain mysterious. Building on structural clues that point to *α*-catenin as the force transducer, we present a quantitative theoretical model for how the catch bond arises, fully accounting for the experimental lifetime distributions. The model allows us to predict the energetic changes induced by tension at the interface between *α*-catenin and F-actin. It also identifies a significant energy barrier due to a network of salt bridges between two conformational states of *α*-catenin. By stabilizing one of these states, this barrier could play a role in how the complex responds to additional *in vivo* binding partners like vinculin. Since significant conformational energy barriers are a common feature of other adhesion systems that exhibit catch bonds, our model can be adapted into a general theoretical framework for integrating structure and function in a variety of force-regulated protein complexes.

## I. INTRODUCTION

The development and maintenance of tissues in multi-cellular organisms requires a diverse array of structural elements that link cells to each other and to the extracellular matrix [1, 2]. For epithelial tissues the main players in cell-cell adhesion are the proteins of the adherens junction complex: transmembrane cadherins and their binding partners that connect the actin cytoskletons of neighboring cells. To understand both healthy tissue architecture and abnormalities that lead to weakening of adhesion in epithelial tumors [3], it is necessary to decipher the underlying molecular mechanisms that regulate the stability of the junctions. Identifying the binding partners of cadherin, their functional roles and interplay under varying environmental conditions, has been a major research goal over the last three decades [2].

The great challenge in achieving this goal is that binding between proteins is not a simple sum of pairwise interactions: the strength of adhesion between any two partners can be allosterically regulated by the presence or absence of other proteins in the complex, as well as conformational changes induced by external factors like mechanical tension [4]. For example, early studies established that the cytoplasmic domain of E-cadherin can bind to *β*-catenin [5, 6], and *β*-catenin can in turn bind to *α*E-catenin [7]. Since the latter was known to independently bind F-actin [8], naively one would assume that αE-catenin would be the bridge linking E-cadherin*β*-catenin to F-actin, forming a minimal recipe for an adherens junction complex (see the schematic model in Fig. 1). However subsequent *in vitro* experiments with purified proteins cast doubts on this model, showing that while E-cadherin/*β*-catenin/*α*E-catenin formed a stable complex, it had significantly lower affinity for F-actin than *α*E-catenin alone [9, 10].

**FIG. 1.**
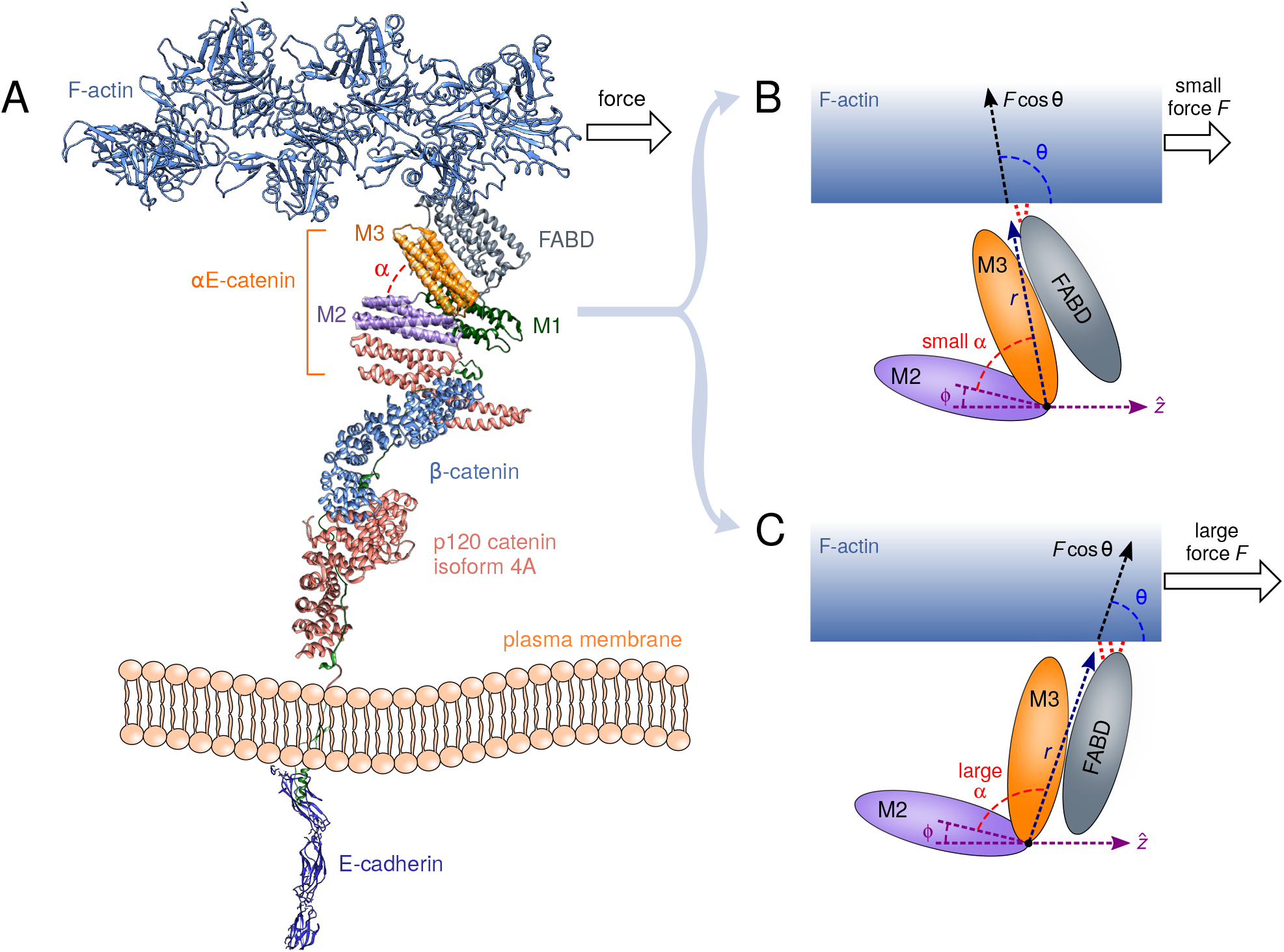
A schematic diagram showing hypothetical conformational changes of the cadherin-catenin-actin complex under force. A) A cartoon of the complex. In the absence of a crystal structure of the entirety, the diagram is drawn from the following PDB structures of various components: 3Q2V (E-cadherin), 3L6X (p120 catenin), 1I7W (*β*-catenin), 4IGG (*α*E-catenin), 1M8Q (F-actin). The arrangement of the structures relative to one another is a guess for the purposes of illustration. The theoretical model described in the text is independent of the details of this arrangement. B) The M region of *α*E-catenin, showing a conformation with small angle *α* between the M2 and M3 domains, favored at lower forces. The interactions (red dashed lines) between the adjacent F-actin binding domain (FABD) and F-actin depend on the conformational state of *α*E-catenin. C) Same as B, but in the large angle conformation, favored at larger forces. This results in an enhancement of FABD-actin interactions, leading to catch bond behavior.

This puzzling result was only clarified three years ago, when Buckley *et al.* added one more ingredient into the mix: applying physiological (pN-level) forces to the entire cadherin-catenin-actin (CCA) system in an optical tweezer [11]. Such external forces mimic the mechanical loads which the complex would feel *in vivo*, and thus would be a more realistic context to study complex formation than the earlier experiments in the absence of load. The results were dramatic: the mean lifetime of the CCA complex increased by a factor of 20 as force was increased from 0 to 10 pN (see Fig. 3), an unusual force-induced strengthening known as catch bonding [12]. The lifetime then fell off exponentially at higher forces, the conventional slip bond decay expected for most biological bonds under tension. The minimal CCA model of the adherens junction gained a new dimension of dynamic complexity: under the right amount of external load, the bond with actin is stabilized up to lifetimes of ~ 1 s, perhaps long enough for vinculin, an additional binding partner between *α*E-catenin and actin to attach and strengthen the junction [13, 14].

Catch bonding has now been observed in a variety of adhesion and receptor proteins complexed with particular ligands, among them selectins [12], integrins [15], bacterial FimH [16], and the *αβ* T-cell receptor [17]. The phenomenon is not limited to protein-ligand complexes, but can occur even in single knotted proteins [18], *α*-helices [19], and force-sensitive functional groups in polymeric materials [20]. One of the most recent observations has been in vinculin [14] binding to actin, where the degree of strengthening under load also depends on the direction of the force. While all these examples highlight the crucial role of tension in regulating interactions, many of them also share the common feature that the structural and energetic details of how this regulation occurs at the molecular level remain largely a mystery. The force spectroscopy experiments that demonstrate protein-ligand catch bonding reveal only the distributions of unbinding times at different forces. We know from very general theoretical considerations that the underlying free energy landscape of a catch bond must necessarily be complex: a simple landscape with a single bound state energy well, and an end-to-end extension that increases monotonically with force, will always yield slip bond behavior [21]. Thus the most likely scenario for catch bonding is a landscape with heterogeneous bound states [22], corresponding to different molecular conformations that can dynamically interconvert under force. But for any specific catch bond system, like CCA, this hypothesis leads to a host of difficult questions: what are the structural differences between the different conformational states? What are the energy barriers between those states? For each state, what are the associated changes in the interaction energies at the bond interface, which are ultimately responsible for the catch bond behavior?

Modeling can assist in tackling these issues, but all current theoretical approaches, despite their various strengths, fall short of being able to directly answer the above questions. The most widely used descriptions of catch bonds are phenomenological [23–27], typically based on a kinetic network of strongly and weakly bound states [24, 28], with force biasing the system toward the strong state. While these models can fit experimental data and capture the essential conceptual basis of catch bonding—conformational heterogeneity—they are expressed in terms of transition rates between states. There is no direct connection between the fitted parameters and the structural features of those states, no way of estimating energy barriers, and no ability to rationalize or predict the results of mutation experiments on the bond lifetimes. Atomistic molecular dynamics simulations give important structural insights [29–32], but have their own limitations: conformational transitions and bond breaking in adhesion complexes at physiological forces typically occur on timescales (ms – s) many orders of magnitude larger than those accessible by allatom simulations, precluding direct comparison to force spectroscopy experiments. Thus a compromise is needed, an approach that is able to fit experimental data, but with results that also have a concrete structural interpretation.

A recent study on the catch bonding in P- and L-selectin adhesion proteins pointed to a possible solution to this problem, introducing a novel, structure-based theory [33]. It provided an analytically solvable model for the mean bond lifetime, whose parameters could be directly linked to the energetics of the interface between the selectin protein and its ligand, as well as structural length scales in the complex. All the fitted parameters were physically reasonable, and in particular the extracted energies were consistent with available crystal structure data on the hydrogen bonding network at the interface. Such a model could for the first time rationalize how particular interfacial energy changes due to mutations would affect the observable bond dynamics. Unfortunately even this approach has an important shortcoming: it assumes the structural transition that occurs under force (in this case the rotation of two selectin domains with respect to each other) does not involve a significant energy barrier. In other words, the transition occurs on timescales much shorter than the mean bond lifetime. At any given force, the model thus yields a probability distribution of lifetimes (also known as a bond survival probability) that is single-exponential.

While the selectin-ligand and other systems [25, 34, 35] considered in Ref. [33] do exhibit single-exponential survival probabilities experimentally, the majority of adhesion systems where data is available do not, including CCA [11, 15, 36–39]. Thus there is a need for a model that is structure-based, analytically tractable, and which can account for the full complexity of bond survival probabilities observed empirically. The theory developed in the current work fulfills all these criteria. It reproduces the experimental lifetime distributions of CCA, and also links them to existing structural information on the conformations of *α*E-catenin. It provides the first estimates of the energy barrier height between these conformations as the complex remodels under force, as well as the resulting energetic changes at the actin interface. These predictions allow us to suggest a future set of experiments to validate the model. They also give insights into the role of the catenin energy barrier in physiological contexts, where a specific conformation of CCA may be required for efficient binding of vinculin to further stabilize the complex [14]. While our focus is on a single system, the theory framework itself is quite general, and can be be readily adapted to other cases. It subsumes earlier models of bond dynamics as special cases in certain limits, including both the barrier-less selectin model and conventional Bell model for slip bonds. It thus has the potential to provide a unified analytical formalism for interpreting data from the entire spectrum of force-regulated adhesion complexes seen in nature.

## THEORY

### Structure-based model

The key structural hypothesis underlying our theory is that conformational changes in the CCA complex induced by force allosterically regulate the interaction strength between F-actin and the C-terminal F-actin binding domain (FABD) [8] of *α*E-catenin (see schematic model in Fig. 1). In the absence of a crystal structure of the FABD-actin interface, many questions remain about its molecular details [40, 41], and among the goals of our approach is to elucidate the overall actin-FABD bond energy and how it varies between different CCA conformations. The precise nature of the conformational changes that occur under tension is also not definitively established, though various lines of evidence point to the central role played by *α*E-catenin as the force transducer [42, 43], including recent dynamic FRET visualization of reversible conformationalchanges in the central domains of *α*E-catenin in a CCA complex under tension in living cells [44]. Fragmentary crystal structures of these central domains [43] suggest the potential of two alpha-helical bundles known as M2 and M3 (residues 396-506 and 507-631 respectively) to adopt different angles with respect to each other. The angle between the bundles (denoted by *α* in Fig. 1) is likely to alter under applied tension, and thus the rotation of M3 with respect to M2 is a natural candidate for the main force-sensitive conformational change [32, 43]. For a catch bond to exist, conformations with small α should be associated with weaker FABD-actin binding, and those with larger *α* with stronger FABD-actin binding. As applied tension biases the system toward the latter conformations, this will lead to a regime where the effective bond lifetime increases with force. This rotation mechanism of catch-bond formation, where the relative orientation between two protein domains is coupled to the bond strength, has proven successful in explaining both experimentally and theoretically the catch bonds in several selectin systems [33, 45], and has recently been suggested as the underlying mechanism in catch bonds between the Notch receptor and certain ligands [36]. One important complication for *α*E-catenin, not present in the selectin cases, is the existence of a significant energy barrier to rotation: crystal structures [41, 43] and molecular dynamics simulations [32] highlight a number of salt bridges among the M-domains that stabilize the small-*α* orientation of M2 and M3. This will prove a crucial ingredient in explaining the dynamics and functional role of the bond, as we will discuss in more detail later.

Synthesizing all these structural considerations into an analytically tractable model, we will posit a minimal Hamiltonian *U*(*r, θ*) for the FABD-actin bond. The conformation-dependence of the bond is encoded in two structural variables (see Fig. 1): i) the magnitude *r* = |r| of the vector r between the rotation pivot point (i.e. the junction of the M2 and M3 domains) and the FABD-actin interface; ii) the angle *θ* between r and the applied force 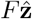 acting on the bond through the actin. The overall geometry of *α*E-catenin relative to actin in Fig. 1 mimics the optical tweezer experimental setup of Ref. [11], whose bond lifetime results we will analyze. That setup was in turn inspired by electron tomographic images showing the organization of actin filaments near the membrane relative to CCA complexes. Fixing 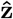 as the actin direction, the M2 domain could have an offset angle *ϕ* relative to z, making the relationship between the M2-M3 domain angle *α* and *θ* have the form: *α = π – θ – ϕ*. Because of steric effects between the domains and the nature of their junction, we assume the angle α can only take on values in some range *α*_min_ ≤ *α* ≤ *α*_max_, which means *θ* is restricted to the corresponding range *θ*_max_ ≥ *θ* ≥ *θ*_min_, where *θ*_max(min)_ ≡ *π* – *α*_min(max)_ – *ϕ*. The Hamiltonian *U(r, θ)* has the form:

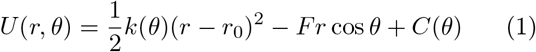

where

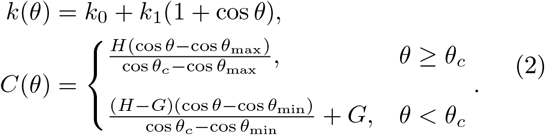

Let us consider each of the terms in Eq. (1) in turn. The first term in the Hamiltonian *U* is an effective bond elastic energy with angle-dependent spring constant *k(θ)* and natural bond length *r*_0_. The distance *r* serves as an effective reaction coordinate for the bond, with bond rupture occurring if *r* > *r*_0_ + *d*, where *d* is the transition state distance. Thus the free energy barrier to bond rupture is *k(θ)d*^2^/2, which depends on the conformation through *k(θ)*. Any angular function *k(θ)* can be expanded in Legendre polynomials *P_l_* (cos *θ*), and for our purposes it is sufficient to keep the two lowest-order terms (*l* = 0,1) in the expansion, *k(θ)* = *k*_0_ + *k*_1_ (1 + cos *θ*), with coefficients *k*_0_, *k*_1_ > 0. This function describes the key feature of the allosteric coupling between the *α*E-catenin conformation and the bond strength: as *θ* decreases under force, *k(θ)* increases, leading to a higher energy barrier to rupture. The extent of the bond strengthening is determined by the magnitude of *k_1_*. In analyzing the bond energetics later, it will be useful to express the role of k_0_, k_1_ equivalently through two energy parameters *E*_0_, *E*_1_ that have simpler physical interpretations. *E*_0_ is the free energy barrier to rupture at *α* = *α*_min_ when *F* = 0, given by *E*_0_ = (*k*_0_ + *k*_1_ (1 + cos *θ*_max_))*d*^2^/2, and *E*_0_+*E*_1_ is the free energy barrier to rupture at *α* = *α*_max_ when *F* = 0. The difference in barrier heights from *α*_min_ to *α*_max_ (responsible for the bond strengthening) is *E*_1_ = *k*_1_(cos *θ*_min_ – cos *θ*_max_)*d*^2^/2.

The second term in *U* describes the coupling of the Hamiltonian to the external applied force of magnitude *F*. It tilts the energy landscape toward larger *r* (increasing the chances of rupture at a given *θ*) and smaller *θ* (or equivalently larger *α*). The final term *C*(*θ*) in *U* describes a free energy barrier between the angular conformational states located at a particular transition angle *α_c_* = *π_c_* – *θ*_c_ – *ϕ*. This effectively subdivides the angular conformational space into two basins: a small interdomain angle region (*α* ≤ *α_c_* or *θ* ≥ *θ*_c_) and a large inter-domain angle region (*α* > *α_c_* or *θ* > *θ_c_*). The barrier passing from small to large *α* has height *H*, and the barrier returning from large to small *α* has height *H* – *G*, with a possible free energy offset *G* between the two basins. As in the case of *k(θ)*, we keep only terms up to linear order in cos *θ*, and make the barrier between the two regions cusp-like for analytical convenience. Using a more complicated form of *C(θ)*, with a smooth rather than cusp-like barrier, would not significantly alter the results of the model (i.e. it would only lead to small corrections ~ *k_B_T* in the fitted results for the energy barriers, where *k_B_* is the Boltzmann constant and *T* the temperature). A representative energy landscape for *U* at *F* = 0 is drawn in Fig. 2 in terms of *r* and *α*, showing the two wells corresponding to the small *α* and large *α* conformational states.

**FIG. 2.**
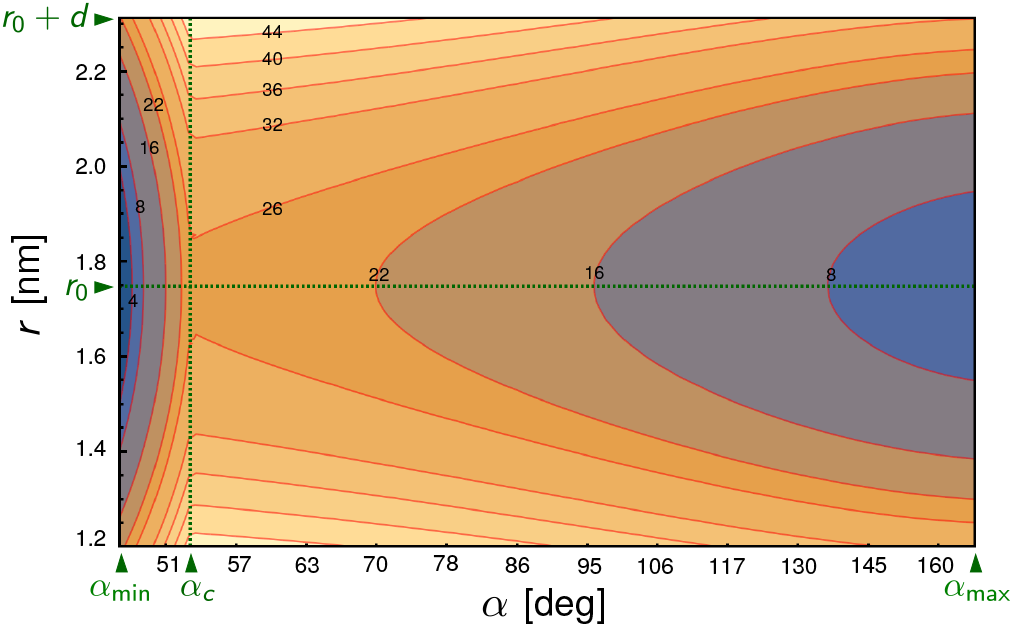
Energy landscape of the Hamiltonian *U* from Eqs. (1)–(2) in terms of *r* and *α* = *π* – *θ* – *ϕ* at force *F* = 0, with the parameters given in Table I and described in the text. Energy contour labels are in units of *k_B_T*. The vertical dashed line corresponds to the transition angle *α_c_*, the horizontal dashed line to the natural bond length *r_0_*, and the top edge to the distance *r*_0_ +*d* beyond which the bond ruptures. The energy barriers to rupture are smaller in the region *α* ≤ *α_c_* on the left, relative to the region *α* > *α_c_* on the right. Since applied force *F* > 0 tilts the landscape toward larger inter-domain angles *α*, the mean bond lifetime will initially increase with force.

The dynamics on this landscape is assumed to be described by diffusion of the vector r obeying a Fokker-Planck equation with potential *U* and diffusivity *D = k_B_T/6πηr*_0_, since the motion corresponds to a rearrangement of a protein domain with characteristic size *r*_0_. Here *η* is the viscosity of water, and for simplicity we ignore any prefactor due to the details of the domain shape in the diffusivity. The corrections due to such a prefactor are small, since it contributes only logarithmically to the fitted energies [33]. Reflecting boundary conditions are assumed at *θ*_min_ and *θ*_max_. The two main dynamical quantities of experimental interest are: (i) the mean bond lifetime *τ*(*F*), defined as the average time it takes to reach bond rupture, *r* = *r*_0_ + *d*, after the onset of an applied force of magnitude *F*. Prior to the force onset, the system is assumed to have equilibrated at zero force, in accordance with the experimental analysis in Ref. [11]; (ii) the survival probability distribution Σ*F*(*t*), defined as the probability that a bond has not yet ruptured by time *t* for a given *F*. The two quantities are related through 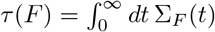

Calculating either *τ*(*F*) or Σ*F*(*t*) analytically is nontrivial for a multi-dimensional potential like *U*, but we can take advantage of the double-well structure of the energy landscape. As shown in detail in the Supplementary Information (SI), we first find approximate analytical expressions for four individual transition rates: crossing the barrier from the small to large *α* well, the reverse transition, bond rupture directly from the small *α* well, and bond rupture directly from the large *α* well. We then combine these expressions into analytical results for *τ*(*F*) and Σ*F*(*t*) in terms of the Hamiltonian parameters.

The final expressions for *τ*(*F*) and Σ_*F*_(*t*) in the SI are rather complex. But as described in the next section, Σ_*F*_(*t*) can be readily incorporated into a maximum likelihood estimation approach to find best-fit Hamiltonian parameters given an experimental data set, i.e. measurements of bond lifetimes at various forces. Moreover *τ*(*F*) reduces to earlier, simpler models of bond dynamics in certain limits. When *H* = *G* = 0, *θ*_min_ = 0, *θ*_max_ = *π*, we exactly recover the expression for *τ*(*F*) in the absence of an angular barrier (and a corresponding Σ*F*(*t*) which is approximately single-exponential), used to describe selectin-ligand catch bonds in Ref. [33] (see details in the SI). If in addition we set *k_i_* = 0, so that *k(θ)* = *k*_0_ becomes independent of *θ*, we do not have any force-enhancement of the bond lifetime. In this limit *τ*(*F*) ∝ exp(–*Fd/k_B_T*), the classic Bell model for conventional slip bonds [46]. The fact that we can smoothly interpolate between different regimes in parameter space, describing qualitatively different modes of force regulation, is one of the strengths of our approach. This allows us, for example, to make predictions for possible mutation experiments that alter the system parameters, and see to what extent the dynamics are robust to such changes.

## III. RESULTS AND DISCUSSION

### Maximum likelihood estimation of the model parameters from force spectroscopy data

To estimate the Hamiltonian parameters and gain insights into the structural mechanisms of catch bonding in the CCA complex, we fit the model to the raw data from the optical tweezer force spectroscopy experiment in Ref. [11]. This data consists of 803 measurements of the bond lifetime under varying force conditions from *F* = 0.7–33 pN, the same dataset whose histogram is depicted in Fig. 4A of Ref. [11]. For a given parameter set and force *F*, the probability to observe a bond lifetime between *t* and *t+dt* is –*dt dΣ_F_(t)/dt*. We could thus construct an overall likelihood function for the data set given the parameters (details in the SI), and maximize it to find the best estimate for the parameters.

For numerical convenience, it was useful to do the fitting in two stages: in the first stage we fixed values for the minimum M2-M3 inter-domain angle *α*_min_ and angle offset *ϕ*, and then maximized the likelihood function over the remaining parameters for these fixed values. In the second stage we then repeated this procedure for different choices of *α*_min_ and *ϕ*, to find the overall optimum. The largest likelihoods occurred in the range *α*_min_ =40 – 50° and *ϕ* = –5 to 5°, yielding results for the remaining parameters identical to within error bars. The best-fit values reported in Table I are for *α*_min_ = 48° and *ϕ* = 0°.

**TABLE I.** Model parameters. Parentheses after the values denote the uncertainty in the last digit.

|  | Physical interpretation | Value |
| --- | --- | --- |
| $E_0$ | Free energy barrier to rupture at small $\alpha$ | 21.8(2) $k_B T$ |
| $E_1$ | Extra free energy barrier to rupture at large $\alpha$ | 5.8(2) $k_B T$ |
| $H$ | Free energy barrier for small-to-large $\alpha$ transition | 25(2) $k_B T$ |
| $G$ | Free energy offset between angular basins | 5(2) $k_B T$ |
| $d$ | Transition state distance | 0.56(1) nm |
| $r_0$ | Distance from hinge to FABD-actin interface | 1.7(2) nm |
| $\alpha_{\min}$ | Minimum angle between M2-M3 domains | 48° |
| $\alpha_c$ | Transition angle between small and large $\alpha$ wells | 53(3)° |
| $\alpha_{\max}$ | Maximum angle between M2-M3 domains | 169(9)° |

The predicted *α*_min_ range is consistent with available structural information. Though the experiment [11] was done using monomeric zebrafish *α*E-catenin, for which there is no crystal structure, we can compare to known homologous structures from other species and computational structure prediction results. 47^°^ was the smallest angle observed in an analysis of available crystal structure fragments of the M2-M3 domains from mouse and human *α*E-catenin [43], and 48° is the M2-M3 angle observed in the individual monomers of the full-length human *α*E-catenin homodimer (PDB: 4IGG) [41]. Plugging the zebrafish *α*E-catenin sequence into the I-TASSER structure prediction server [47, 48] yields an M2-M3 angle of 45 ± 1° among the five best structures.

The theoretical mean bond lifetime *τ*(*F*) is compared to the experimental results from Ref. [11] in Fig. 3, and the analogous comparison for the survival probabilities *Σ*_F_(*t*) at different *F* is shown in Fig. 4. The agreement between theory and experiment is excellent, with the model capturing not only the catch bond trend in *τ*(*F*), but also the clear double-exponential behavior in Σ_*F*_(*t*). As we will discuss in more detail below, the observation of two exponential regimes is closely connected to the presence of a significant energy barrier between the small *α* and large *α* conformations.

**FIG. 3.**
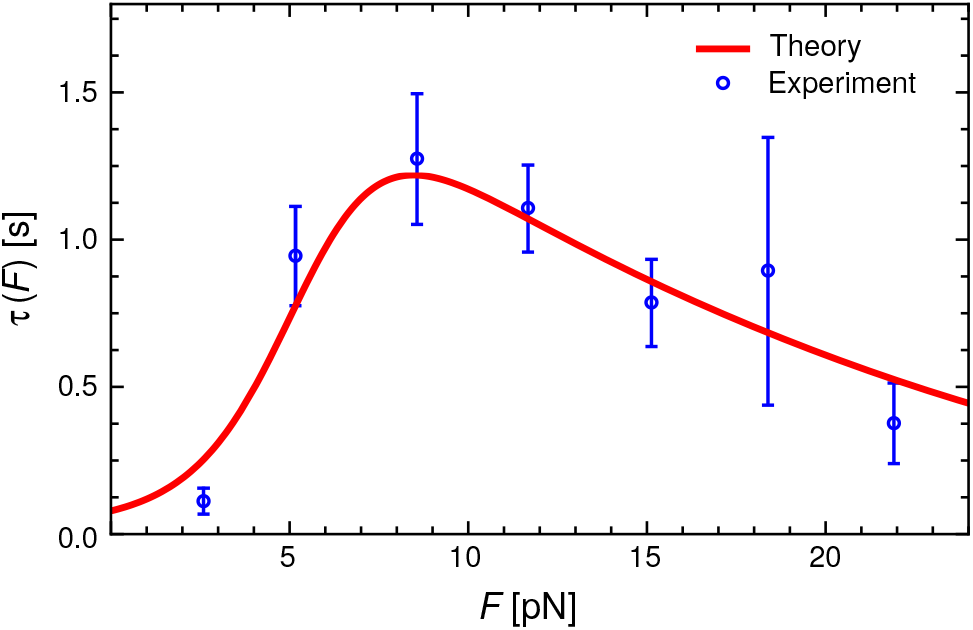
Experimental mean bond lifetime *τ*(*F*) versus force *F* (symbols) from Ref. [11] compared to the theoretical model with best-fit parameters from Table I (curve).

**FIG. 4.**
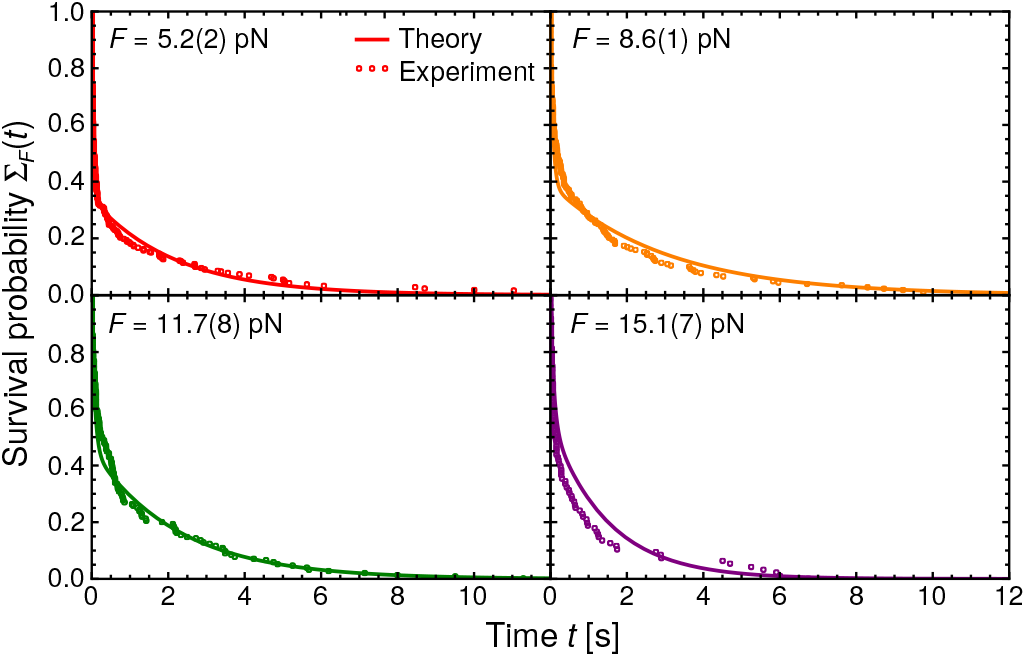
Bond survival probability Σ_*F*_(*t*) versus time *t* for four different forces *F*. Theory results are shown as curves, and the corresponding experimental data [11] as symbols.

### Interpretation of the model parameters, and corroboration from structural data

The value of the model comes not just from the fact that it can fit the experimental data, but that its parameters have a direct physical interpretation that illuminates the structural mechanism of the CCA catch bond. The energy barrier at the transition angle *α_c_* = 53° divides the parameter space into two basins: a narrow basin between *α*_min_ = 48° to *α_c_*, and a much wider basin between *α_c_* and *α*_max_ = 169°. The narrow range suggests the M3 domain is held rigidly in place relative to M2 in the small *α* case, with limited rotational mobility, but once the stabilizing interactions at the hinge between M2 and M3 are broken, M3 can swing out to a larger angle. Of course the idea of solid body rotation about a hinge is a simplification: the protein domains are plastic objects that can continuously deform under tension, but picturing an overall rotation is still a useful first approximation. The parameter *r*_0_ = 1.7 nm, in the simple picture the distance between the hinge and the FABD-actin interface, can more accurately be interpreted as the effective size of the protein regions undergoing reorientation under force.

The strength of the interactions in the hinge region is reflected in the angular energy barrier height *H* = 25 *k_B_T*, whose full significance we will explore below. The existence of this barrier is supported by corroborating evidence from a crystal structure [41] of *α*E-catenin (PDB: 4IGG), which shows five inter-domain salt bridges in the hinge region where the M1, M2, and M3 domains meet (Fig. 5). If each salt bridge roughly contributes 4 – 8 *k_B_ T* to the overall barrier [49], this is consistent with the magnitude of *H*. Molecular dynamics simulations also point to the stabilizing role of the salt bridges. Li *et al.* [32] compared trajectories measuring the M2-M3 angle for the wild-type structure, initially starting in the small *α* state, to trajectories of mutants where one of the salt bridges is disrupted (i.e. E521A or R551A). The latter show the system venturing more readily to larger angles relative to the wild-type, as expected for a smaller barrier *H*.

**FIG. 5.**
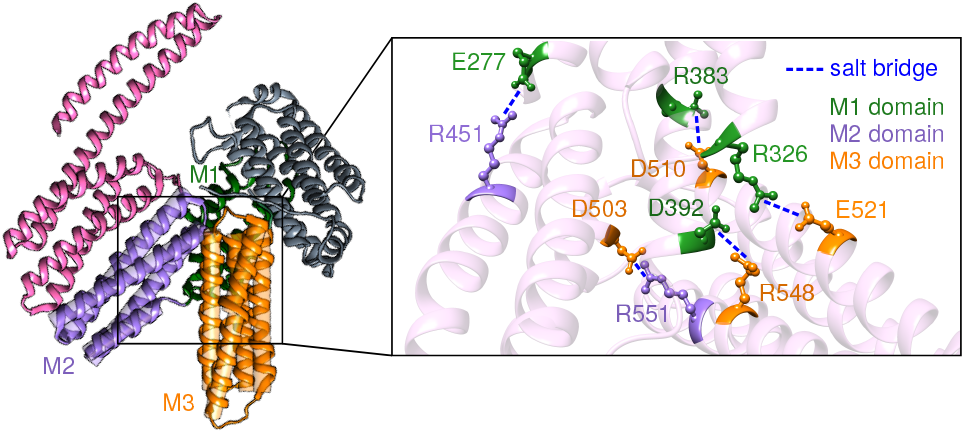
The salt bridge network in the hinge region between the M1, M2, and M3 domains of *α*E-catenin (PDB: 4IGG) [41].

Having two conformational states at small and large *α* in itself does not guarantee catch bond behavior. What leads to lifetime enhancement under force is the fact that these states are allosterically coupled to the strength of the FABD-actin bond, which changes from *E*_0_ = 21.8 *k_B_T* at small *α* to *E_0_+E_1_* = 27.6 *k_B_T* at large *α*. Though we do not have any crystal structure of the FABD-actin interface, it is instructive to compare the value of *E_0_ + E_1_* to a different catch bond system: the P-selectin complex with the ligand PSGL-1, where *E_0_ + E_1_* = 27 *k_B_T* [33] in the extended state favored at larger forces. The peak bond lifetime in P-selectin/PSGL-1 (~ 1.1 s) is also very similar to CCA (~ 1.1 s in Fig. 3). Conveniently we do have the crystal structure of P-selectin-PSGL-1 in the extended conformation (PDB: 1G1S) [50], showing that 20 hydrogen bonds contribute to *E_0_ + E_1_*, consistent with a contribution of 1.2 — 1.5 *k_B_T* per hydrogen bond, typical for hydrogen bonds in proteins [51]. We thus predict a similar number of hydrogen bonds at the FABD-actin interface in the large angle state (or fewer if salt bridges are involved). The allosteric change between the angular states translates into an interface energy difference of *E*_l_ = 5.8 *k_B_T*, about 4-5 hydrogen bonds or one salt bridge.

The energy offset parameter *G* = 5 *k_B_T* plays the important role of biasing the system toward small *α* when the force is small. The equilibrium probability 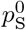 of having *α* < *α_c_* at *F* = 0 is 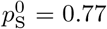 (see SI for the derivation). As *F* is increased, the energy landscape is tilted toward higher *α*, and the barrier to FABD-actin bond rupture shifts from *E*_0_ to *E*_0_ + *E*_l_, causing the lifetime enhancement. But the fact that the system is equilibrated at *F* = 0 before the application of force means that both large and small angle conformations are initially populated. The significant angular barrier *H* and the finite bond lifetime means that these populations do not necessarily have a chance to fully re-equilibrate once *F* > 0 is applied, during the time before rupture occurs.

These two populations, one with a smaller barrier to rupture than the other, explain the distinct double exponential behavior of Σ_F_(*t)* that exists even at the highest forces investigated in the experiment (Fig. 4). To understand this more concretely, a useful quantity is the probability of being in the small *α* state at the moment of rupture, the so-called splitting probability *π*_s_ (details given in the SI). In the hypothetical scenario of arbitrarily long-lived bonds, where there is time for many transitions between the small and large *α* states, *π*_s_ ≈ *p*_s_, the equilibrium probability of being in the small *α* state. But in many cases the bond lifetime is too short for equilibration, and *π*_s_ may be very different from *p_s_*. For example at *F* = 15.1 pN (the last panel in Fig. 4), *p*_s_ = 10^-4^, but *π*_s_ = 0.47. The tiny value of *p*_s_ means that, given enough time, the initial fraction, 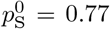, of systems that start at small α should eventually transition to the large *α* state preferred at high forces, and almost never return. If that were actually the case, the survival probability at *F* = 15.1 pN would have been to very good approximation a single exponential, since rupture would occur almost entirely from the large *α* state. In reality, because of the barrier *H* slowing down angular transitions, the majority of those small *α* systems do not have enough time to transition. They thus stay in the small *α* state until rupture, giving a sizable *π*_s_. This leads to a short lifetime exponential regime in Π*F*(*t*), in addition to the longer lifetime exponential regime corresponding to ruptures from large *α*.

The final parameter in the model, the transition state distance *d* = 0.56 nm, represents how much the FABD-actin bond interface can be deformed before rupture. The value is within the range expected of most proteins (< 2 nm) [52]. Putting everything together, we thus see that the fitted model parameters are all within physically realistic ranges, and consistent with all the available evidence both from the Buckley *et al.* experiment and earlier studies.

### Mutations to the angular barrier *H*, and its potential biological role

Disrupting the stability of the hinge region (Fig. 5) with mutations at the M2-M3 inter-face (R551A) or M1-M3 interface (E521A) has been experimentally investigated to probe the role of the hinge in vinculin binding [43]. The underlying presumption is that the large *α* conformation, which is more accessible when the hinge is destabilized, exposes the vinculin binding site in the M1 domain. This would explain the enhanced binding affinity of the R551A and E521A mutants to the D1 domain of vinculin seen in the experiments. Of course in nature, access to the large *α* conformation is controlled not by mutations to the hinge, but by application of force, leading to the speculation that the *α*E-catenin system acts like a force-dependent “switch” [43], with tension favoring a large *α* conformation, which in turn enhances both vinculin and F-actin bond strengths.

In the context of the model, there are two scenarios for what might occur when the salt-bridge network at the hinge is disrupted: (i) the angular barrier energy *H* is decreased, since this is the parameter most directly related to the stability of the hinge, but other parameters in the model remain unaffected; (ii) the decrease of *H* is allosterically coupled to changes in the FABD-actin interfaces energies *E*_0_, *E*_1_ or other structural parameters. The latter would be reminiscent of the case of L-selectin, where experimental mutations at the hinge between the lectin and EGF domains [25] led to allosteric changes in energies at the ligand-binding interface [33]. The possibility of scenario (ii) will have to await future experimental data, but we can explore scenario (i) theoretically. This also allows us to investigate the biological significance of the angular barrier *H*.

Fig. 6A shows what happens to the mean bond lifetime *τ*(*F*) when *H* is decreased from its wild-type value of 25 *k_B_ T* in increments of 5 *k_B_ T* (roughly corresponding to removal of individual salt bridges), while all other parameters are fixed at their Table 1 values. The catch bond behavior is preserved, but with opposite trends at small and large forces: at small forces *τ*(*F*) generally decreases with decreasing *H*, while at larger forces it initially increases by about a factor of two at the maximum, and then decreases gradually. These changes are due to the fact that transitions between the small and large *α* states become easier with decreasing barrier heights. At smaller forces, where the weaker small *α* states are preferred, some fraction of systems that would have ruptured from the stronger large *α* state can now transition to small *α* before rupturing. The converse is true at larger forces, where we now allow more small *α* states to transition to the preferred large α state before rupture.

**FIG. 6.**
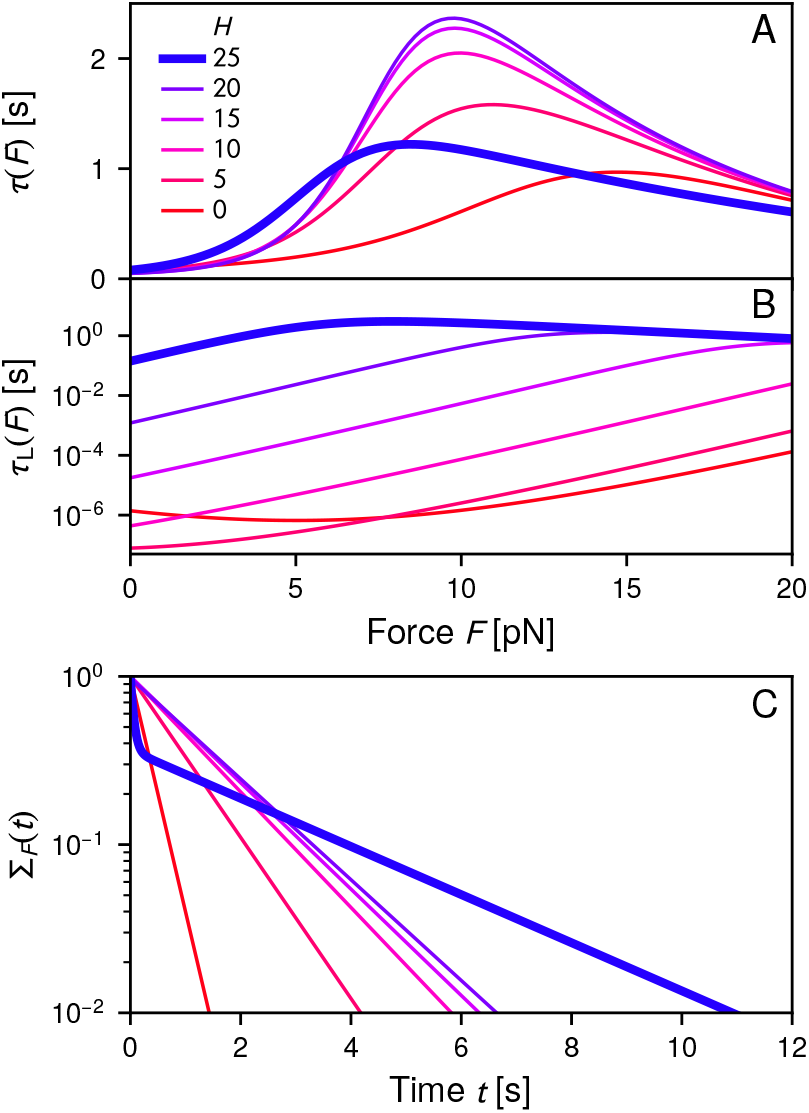
The effects of mutating the angular barrier height *H* from the original value of 25 *k_B_T* down to zero, in increments of 5 *k_B_ T*, leaving all other model parameters fixed at their Table 1 values: A) the mean bond lifetime *τ*(*F*); B) the mean lifetime *τ_L_* (*F*) of remaining in the large angle conformational state, *α > α_c_*, measured from the initial time of entry into the state; C) the survival probability Σ*_F_* (*t*).

Consistent with this, the lifetimes within each angular domain are drastically affected by the mutation. Fig. 6B shows *τ*_L_(*F*), the mean duration of the large *α* state (from initial entry into the state until either rupture occurs or a transition to small *α*; see SI for details). For the wild-type value *H* = 25 *k_B_ T*, there is a broad force region, *F* ≈ 4 – 18 pN, where the large angle state survives for macroscopic times comparable to the maximum bond lifetime, *τ*_L_(*F*) > 1 s. When *H* = 20 *k_B_T* this region is decreased to *F* ≈ 12 – 18 pN, and then vanishes entirely at smaller *H*. With a decreasing barrier, the time spent at large α becomes significantly briefer, reduced by 4-6 orders of magnitude at *H* = 0. At *H* = 25 *k_B_ T* a typical system trajectory may have involved zero or one transition across the angular barrier, and then rupture. In contrast at smaller *H* the system makes a large number of angular transitions before the bond breaks. The result is that the double-well nature of the energy landscape is averaged out, and the survival probability Σ*_F_*(*t*) switches from double-exponential at *H* = 25 *k_B_ T* to mainly single-exponential at *H* ≤ 20 *k_B_T*, as seen in Fig. 6C.

Thus while the presence of a large *H* barrier is not necessary for catch bonding, it is necessary to stabilize the large *α* conformational state so that it persists for long durations. A larger *τ*_L_(*F*) over a wide force range comes at the price of a somewhat smaller maximum *τ*(*F*). But this may be biologically preferred if the macroscopic duration of the large *α* state is necessary to allow time for additional binding partners (like vinculin) to dock before rupture or the transition to small *α*. Indeed two potentially fruitful future lines of experimental inquiry would be: a) to first study the CCA catch bond under different mutations to the *α*E-catenin hinge region. The mutations would have a clear signature of their effect on *H* by the change in the nature of the survival probability distribution [Fig. 6C]. Whether the response of *τ*(*F*) would follow the trend in Fig. 6A would determine if scenario (i) were true, or whether additional allosteric effects like in scenario (ii) are also present; b) to study the binding affinity or bond lifetime of vinculin to the CCA complex under these same mutations. This would elucidate whether the increased lifetime of the large *α* state, facilitated by the angular barrier, is also required for effective vinculin binding. One can also imagine an alternative vinculin binding mechanism like induced fit, where its affinity might be independent of the lifetimes or relative populations of the *α*E-catenin conformational states.

## IV. CONCLUSIONS

The model presented here is the first quantitative, structural model for the catch bond in the cadherin-catenin-actin complex. It provides a full interpretation of the force spectroscopy data from the Buckley *et al.* experiment [11], highlighting the central role of *α*E-catenin as a force-transducing conformational switch [42–44]. The switch mechanism, based on small and large angle catenin conformations with different FABD-actin bond strengths, is to date the most plausible molecular explanation of the CCA catch bond. Force induces a small-to-large angle transition over a substantial energy barrier resulting from a network of salt bridges. This energy barrier, captured in the parameter *H* in our model, leads to the double-exponential survival probabilities seen experimentally. Additionally, once the system transitions to the large α conformation, the barrier allows it to remain there a significant fraction of the bond lifetime, perhaps facilitating the binding of other proteins like vinculin which play major roles in the physiological complex. While the model parameters are consistent with all the available evidence, including structural information about the *α*E-catenin hinge region, full corroboration of the mechanism will require further experiments to check whether alterations in the *α*E-catenin conformational stability have the posited effects on bond observables. Moreover, future crystal structures of the FABD-actin interface would allow verification of the *E*_0_ and *E*_0_ + *E*_i_ energy scales predicted by our approach.

Of course it is always possible that an alternative conformational mechanism will emerge for the CCA catch bond. Any competing explanation will still have to include a conformational change whose dynamics are slowed down by an energy barrier ≫ *k_B_ T*, since this is the only way to have a catch bond with double-exponential survival probabilities. One of the attractive features of our model is that it can be readily adapted for such an eventuality. The current Hamiltonian is expressed in terms of bond distance and inter-domain angle, but analogous Hamiltonians can be formulated, replacing the angle with another conformational coordinate. The model can even generalize to more than two conformational basins in the energy landscape, separated by different barriers, if the structural evidence points in that direction. The basic approach stays the same, and analytical expressions for the bond lifetimes and distributions can always be derived to fit to experimental data. Given the ubiquity of multi-exponential lifetime distributions in catch bonding systems [11, 15, 36–39], implicating conformational transitions with non-trivial energy barriers, our approach thus might provide a universal framework for structural modeling of catch bonding. And it is not only limited to multi-exponential distributions, since single-exponential behaviors (for both catch and slip bonding) are just special cases of the model parameters. The usefulness of our theory starts at the cadherin-catenin-actin system, but hopefully will not end there.

## ACKNOWLEDGMENTS

The authors thank A. Dunn for sharing the experimental force spectroscopy data. S.A. thanks A. Dunn, D. Huang, and W. Weis for useful discussions. This work was supported by the National Science Foundation (CAREER BIO/MCB 1651560). Computational resources were provided by the Case Western Reserve University High Performance Computing Cluster.

**Supporting Information for “Unraveling the mechanism of the cadherin-catenin-actin catch bond”**

Shishir Adhikari, Jacob Moran, Christopher Weddle, and Michael Hinczewski

Department of Physics, Case Western Reserve University, Cleveland OH, 44-106, U.S.A.

## I. DERIVATION OF MEAN LIFETIME *τ*(*F*) AND SURVIVAL PROBABILITY Σ*_F_*(*t*)

The following sections contain a full derivation of the main observable quantities of interest, the mean bond lifetime *τ*(*F*) and survival probability Σ_F_(*t*). Since the derivation involves a large number of individual components, Table S1 summarizes the main analytical quantities, their meaning, and the equations where they are defined.

**TABLE S1.** Summary of main analytical results in the Supporting Information, with corresponding equation numbers.

| Quantity | Meaning | Equation(s) |
| --- | --- | --- |
| $U(r, \theta)$ | bond Hamiltonian | (S1)-(S2) |
| $V(r, \theta)$ | effective Fokker-Planck potential energy | (S6) |
| $k_{10}$ | transition rate from state 1 to 0 (rupture) | (S15)-(S17) |
| $k_{20}$ | transition rate from state 2 to 0 (rupture) | (S18)-(S19) |
| $k_{12}$ | transition rate from state 1 to 2 | (S26)-(S27) |
| $k_{21}$ | transition rate from state 2 to 1 | (S28) |
| $\tau_i$ | mean first passage time from state $i$ to rupture | (S35) |
| $p_i^0$ | probability of state $i$ at time $t = 0$ | (S36) |
| $\tau$ | mean bond lifetime | (S37) |
| $\Sigma_F$ | survival probability | (S38)-(S39) |
| $\tau_L$ | mean duration of large $\alpha$ conformation (state 2) | (S42) |

### A. Fokker-Planck equation describing the bond dynamics

The theoretical model of the bond dynamics is based on diffusion of the bond vector r = (*r, θ, ϕ*) on an energy landscape defined by the Hamiltonian *U(r, θ)* in Eqs. 1–2 in the main text:

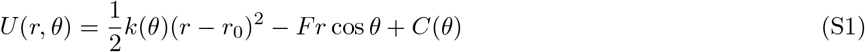

where

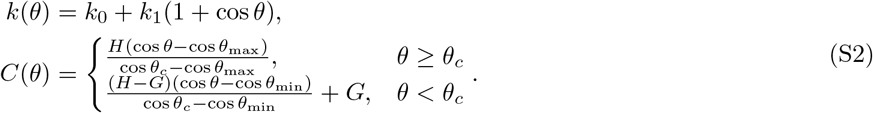

To restrict the dynamics to the angular region *θ*_min_ ≤ θ ≤ *θ*_max_, we assume *U(r, θ)* = ∞ for *θ < θ*_min_ and *θ* > *θ*_max_. Note that *U(r, θ)* depends on the applied force *F* on the system, so every observable derived from *U(r, θ)* below also implicitly depends on *F*, even if the dependence is not explicitly indicated in the notation.

Given a diffusivity *D* = *k_B_T*/6*πηr*_0_, the probability Ψ(r,*t*) to find the system with vector r at time *t* obeys a Fokker-Planck equation in spherical coordinates of the form:

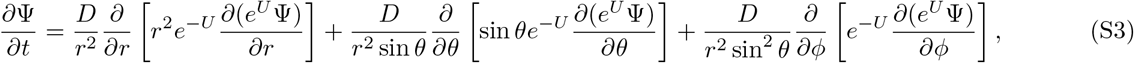

Note that throughout the supporting information we will work in units where *β* = (*k_B_T*) ^-1^ = 1, so that all energies are effectively measured in units of *k_B_T*. Since *U(r, θ)* is independent of *ϕ*, we can define a marginal probability *P(r, θ, t)* by multiplying Ψ with the spherical Jacobian and integrating over the angle *ϕ*,

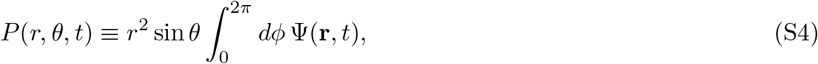

allowing us to write Eq. S3 as a 2D Fokker-Planck equation in terms of *P(r, θ, t)*,

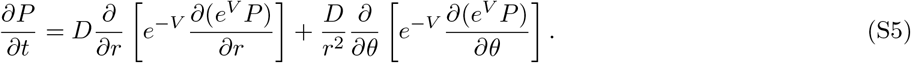

Here *V(r, θ)* is an effective potential defined by

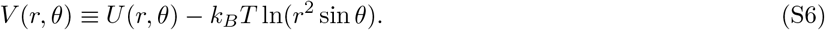

We will not be interested in solving Eq. (S5) directly, but rather answering a closely related question: the mean first passage time (MFPT) to escape from a region in parameter space. Consider a region *R* of the (*r, θ*) space, with boundary *∂R*, and let us focus on some subset 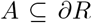 of this boundary. Let *τ_RA_ (r, θ)* denote the MFPT to any point on *A*, given that we started at some point (*r, θ*) in the interior of *R* at *t* = 0. We assume we have chosen *R* such that there are reflecting boundaries on the portion of *∂R* not in *A*, namely *U(r, θ)* = ∞for (*r, θ*) ∈ *∂R \ A*. This guarantees that *τ_RA_ (r, θ)* is finite, and satisfies the backward Fokker-Planck equation [1],

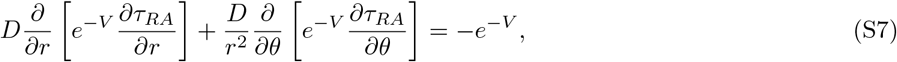

with absorbing boundary conditions *τ_RA_ (r, θ)* = 0 for (*r, θ*) ∈ *A*. To directly solve for the main experimental observable of interest, the mean bond lifetime, we would set *R* to be the entire parameter space region where the bond is intact, *r < r_0_ + d* ≡ *b, θ*_min_ ≤ *α* ≤ *α*_max_, and set the absorbing boundary *A* to be the line *r = b, θ*_min_ ≤ *θ* ≤ *θ*_max_. Unfortunately, given the complicated form of the energy landscape, Eq. (S7) does not easily lend itself to an analytical solution for this choice of *R* and *A*. We will work around this problem by describing the escape dynamics from smaller portions of the parameter space, where Eq. (S7) is more amenable to approximation, and then piece together the various results to get a good estimate of the mean bond lifetime. This same approximate piece-wise approach will also yield the survival probability.

### B. Partitioning the parameter space into conformational regions

**FIG. S1.**
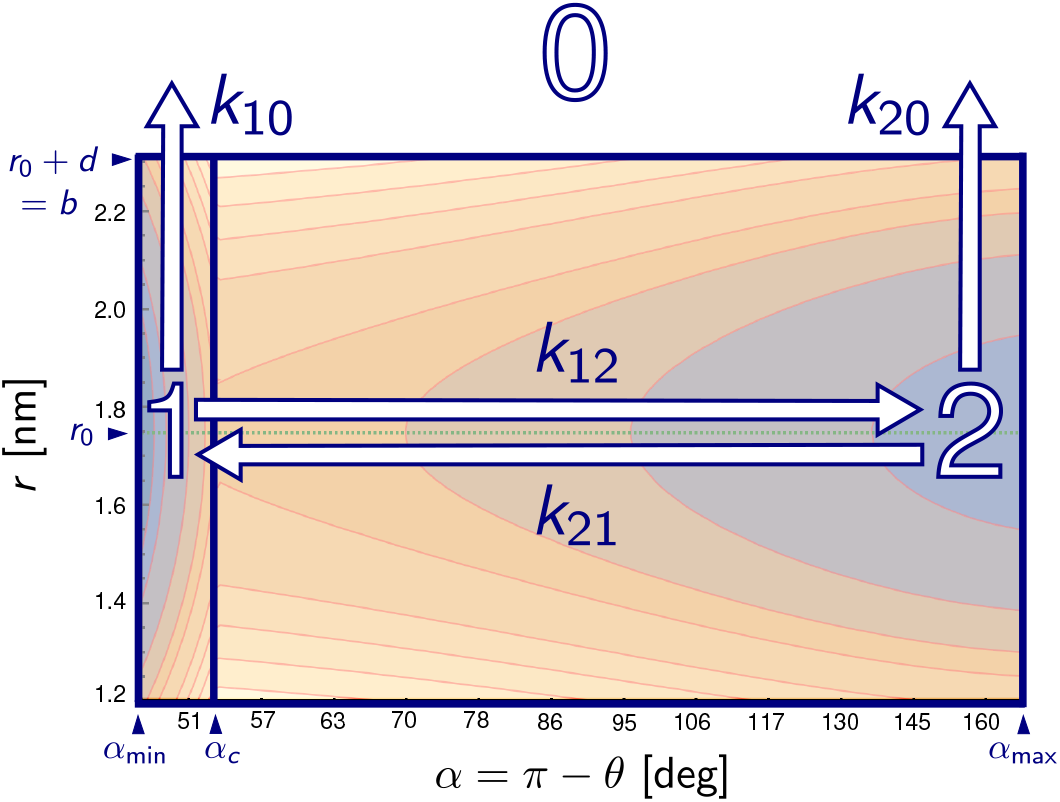
The energy landscape from Fig. 2 of the main text, partioned into regions of the (*r, α*) parameter space that reflect different conformational states: state 0 corresponding to the bond ruptured, state 1 corresponding to the bond intact with angle *α*_min_ ≤ *α* ≤ *α_c_*, and state 2 corresponding to the bond intact with angle *α_c_* ≤ *α* ≤ *α*_max_. The arrows depict the transition rates *k*_10_, *k*_20_, *k*_12_, and *k*_21_ between the various states, described in the text.

The energy landscape of Eqs. (S1)–(S2) allows us to partition the (*r, θ*) parameter space, or equivalently the space of (*r, α = π – θ*), into domains representing different conformational states, as illustrated in Fig. S1. The region where the bond is intact (*r < b*) and the angle α is small (*α*_min_ ≤ *α* ≤ *α_c_*) is denoted as state 1, the corresponding region with an intact bond and large angle (*α_c_* ≤ *α* ≤ *α*_max_) is denoted as state 2, and the region where the bond is ruptured (*r ≥ b*) as state 0. If the bond is intact at time *t* = 0, the dynamics of the system will consist of diffusion on the energy landscape, possibly making a number of transitions between states 1 and 2, before eventually the *r = b* boundary is crossed and bond rupture occurs upon entry to state 0.

The diffusive dynamics exhibit a separation of time scales: the time to equilibrate within each state is typically 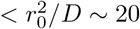 for the biologically relevant parameter ranges we consider, which is orders of magnitude smaller than the typical times to escape each state (which can be as large as ~ 1s for the energy barriers in our case). We can thus assume that upon entering either state 1 or state 2, the system will rapidly be driven toward the bottom of the corresponding energy well, and spend considerable amounts of time in the vicinity of the local energy minimum, with brief excursions up the slopes of the well (eventually one of which will carry it into the bond rupture region, or a transition to the other angular state).

Given this partitioning of the parameter space, we can define escape rates from the different states through different boundaries, related to the reciprocal of the escape MFPT introduced in the previous section. These rates (*k*_10_, *k*_20_, *k*_11_, and *k*_21_) are depicted as transition arrows in Fig. S1. Let *k_10_* be the probability per unit time to escape state 1 to state 0 through the bond rupture (*r = b*) boundary, conditioned on not passing through state 2. We set *k*_10_ = 1/*τ*_10_(*r_1_, θ_1_*), where *τ*_10_ is the solution to Eq. (S7) with the region *R* corresponding to state 1, an absorbing boundary *A* at the border with state 0 (*r = b*) and a reflecting boundary condition imposed at the border with state 2 (*α = α_c_*). The starting position (*r_1_, θ_1_*) is the position of the local minimum of the effective potential *V(r, θ)* in state 1. Using any other starting position in the vicinity will not substantially alter the result, given the fast equilibration time in the well. We define the escape rate from state 2 to state 0, conditioned on not passing through state 1, as *k*_20_ = 1/*τ*_20_(*r_2_, θ_2_*), with *R* being state 2, an absorbing boundary *A* at the border to state 0, and a reflecting boundary at the border to state 1. The position (*r_2_, θ_2_*) corresponds to the local minimum of *V(r, θ)* in state 2. The interwell transition rates *k*_12_ and *k*_21_ are defined analogously: *k*_12_ = 1/*τ*_12_(*r*_1_, *θ*_1_) is the escape rate from state 1 to state 2, conditioned on the bond not rupturing (reflecting boundary at *r = b*), and *k*_21_ = 1/*τ*_21_(*r_2_, θ_2_*) is the reverse rate from state 2 to state 1.

In the next two sections (1.3 and 1.4) we will find approximate expressions for *k*_10_, *k*_20_, *k*_12_ and *k*_21_ from Eq. (S7), and then in section 1.5 we will show how we can put these together to get analytical results for the mean bond lifetime *τ(F)* and survival probability Σ*_F_(t)*.

### C. Deriving expressions for *k*_10_ and *k*_20_

To find an expression for *k*_10_ = 1/*τ*_10_(*r_1_, θ_1_*), we note that *τ*_10_ satisfies Eq. (S7) with the region *R* corresponding to *r < b, α_min_ ≤ α < α_c_*. The *α* angular range is equivalent to *θ_c_* μ*θ* ≤ *θ*_max_. We impose reflecting boundary conditions at *θ_c_* and *θ*_max_, and there is a natural reflecting boundary condition at *r* = 0 because of the logarithmic term in the definition of *V(r, θ)* in Eq. (S6). The absorbing boundary *A* is *r = b*, the border with state 0. We will reduce the dimensionality of the problem by integrating both sides of Eq. S7 over the *θ* range of *R*,

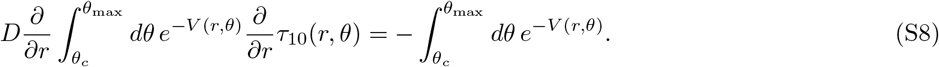

The second term on the left hand side in Eq. S7 vanishes after the integration because exp(-*V(r, θ)*) = 0 at *θ* = *θ_c_* and *θ* = *θ*_max_ because of the reflecting boundary conditions.

Because of the *e^−V(r,θ)^* terms inside the integrals on both sides of Eq. (S8), the dominant contribution to the integrals at any particular value of *r* occurs when *V(r, θ)* reaches a minimum with respect to *θ* inside the state 1 range *θ_c_* μ*θ* ≤ *θ*_max_. This happens at some value of *θ* = *θ*_m1_(*r*) for a given *r*. Thus we can treat Eq. (S8) as an equation for *τ*_10_(*r, θ_m1_(r)*), which we will write in condensed notation as just *τ_10_(r)*. Note that we are interested in *τ*_10_(*r*_1_), since *α*_1_ = *α*_m1_(*r*_1_) is just the position of the well minimum at *r*_1_. In this approximation Eq. (S8) becomes

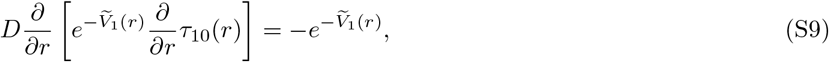

where the effective 1D potential 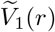 is given by

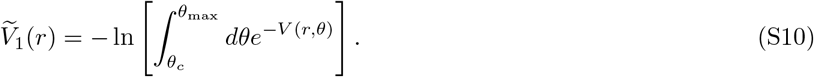

With the absorbing boundary condition *τ*_10_(b) = 0, Eq. S9 can be solved for *τ*_10_(*r*),

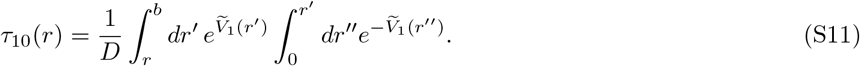

The function 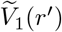 is a monotonically increasing function of *r’* at large *r’*. Due to the presence of the 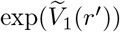 term, the integral over *r’* in Eq. S11 gets its dominant contribution from *r’* near the upper limit b. Conversely because of the 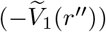 term, the integral over *r"* gets its dominant contribution near 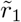, the position where 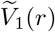 reaches a minimum. To simplify Eq. (S11), we thus make two approximations: (i) expand 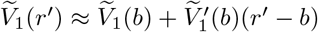; (ii) assume 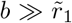, so the upper limit in the integral over *r"* can be replaced by ∞. Along similar lines, if the starting position is *r = r*_1_, the precise value of the lower limit on the *r’* integral has a negligible effect on the result, so we can replace it with 0. Eq. (S11) can then be evaluated to yield an expression for *τ*_10_(*r*_i_),

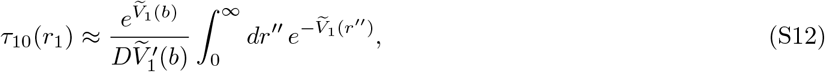

where we have kept the largest contributions to the result. The right-hand side does not have a dependence on the value of the starting position *r*_1_, consistent with the assumption of fast equilibration within the well. We denote the integral in Eq. S12 as

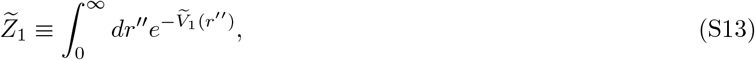

which needs to be evaluated to get a closed form expression for *τ*_10_(*r*_1_). Because this cannot be done exactly, we will use a saddle-point approximation by expanding 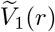 around its minimum at 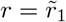,

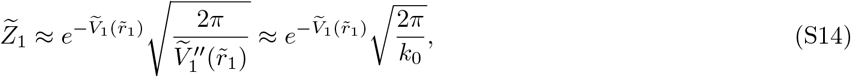

where we have used the fact that 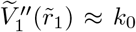 in the limit of large *k*_0_ (which is valid when the barrier to rupture *E*_0_ ≫ *k_B_T*, since *E*_0_ = (*k*_0_ + *k*_1_(1 + cos *θ*_max_))*d*^2^/2).

Putting everything together from Eqs. (S12)–(S14), we can find *τ*_10_(*r*_1_) explicitly. This requires carrying out 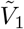 integrals using the definition of Eq. (S10), and approximating 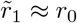. This leads to a final expression for *k*_10_:

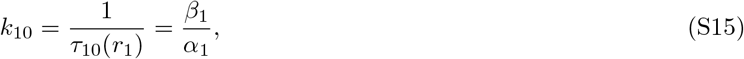

where

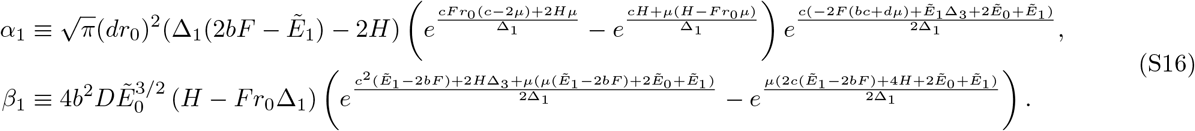

In the above expressions, as well as the ones for the other rates below, we will use a set of abbreviated notations, as follows:

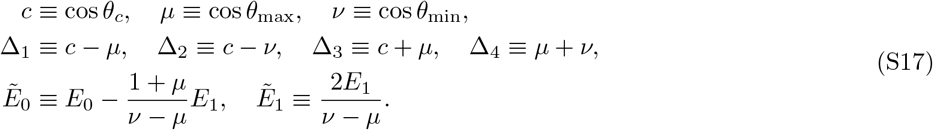

For *k*_20_ = 1/*τ*_20_(*r*_2_, *θ*_2_), the derivation proceeds exactly analogously to the one for *k*_10_, except that the region *R* now corresponds to *r* < *b, θ*_min_ ≤ *θ* ≤ *θ_c_*. The final expression for *k*_20_ is:

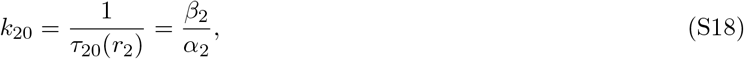

where

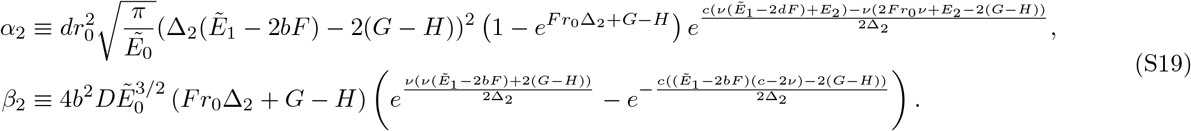

### D. Deriving expressions for *k*_12_ and *k*_21_

Let us first consider the transition rate *k*_12_ = 1/*τ*_12_(*r*_1_,*θ*_1_) from state 1 to state 2, conditioned on the bond not rupturing. The starting point (*r_1_,θ_1_*) is at the local energy minimum in state 1, and we will place the absorbing boundary at some angle *θ* < *θ_c_*, beyond the angular energy barrier at *θ_c_* that defines the border with state 2. Once we are not in the immediate vicinity of the barrier top, the precise location of the absorbing boundary within state 2 does not significantly change the value of *τ*_12_. This is because once the system has overcome the barrier to transition from state 1 to state 2, it rapidly descends into the state 2 energy well. Using this freedom, we will choose the absorbing boundary *A at θ = θ*_2_, the position of the local energy minimum in the state 2 well.

We choose the region *R* to have an *r* range between 0 and *b*, with a reflecting boundary imposed at *r = b*. The logarithmic term in *V(r, θ)* in Eq. (S6) provides another reflecting boundary at *r* = 0. Integrating both sides of Eq. S7 over this *r* range gives:

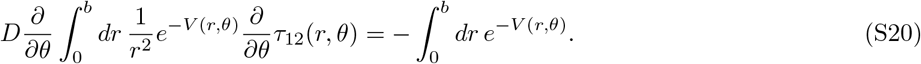

The first term in Eq. S7 vanishes under integration because of the reecting boundary conditions.
The dominant contribution to the integral on the left-hand side of Eq. S20 for a given angle *θ* occurs at *r = r_m_(θ)*, where *V (r, θ)* is minimal with respect to *r* at that *θ*. To a good approximation *r_m_(θ) ≈ r_0_* for the force and parameter ranges we consider. We can thus treat Eq. (S20) as an effective equation for *τ*_12_(*r_0_, θ*), which we will denote compactly as *τ*_12_(*θ*). We are ultimately interested in getting an expression for *k_12_* = 1/*τ*_12_(*θ_1_*). Using this approximation, we can rewrite Eq. (S20) as:

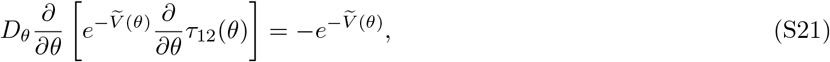

where 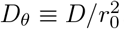, and the effective 1D potential 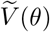 is given by

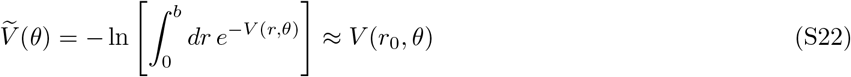

In the second expression on the right, we have kept only the most significant contribution from the saddle-point
approximation of the integral.

With the absorbing boundary condition *τ*_12_(*θ*_2_) = 0, Eq. S21 can be solved for *τ*_12_(*θ*):

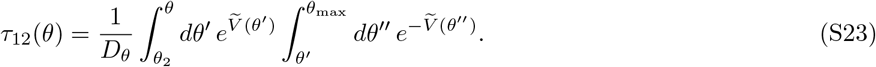

Substituting 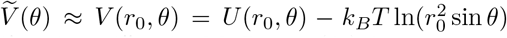 allows us to rewrite Eq. (S23) in terms of integration variables cos *θ’* and cos *θ"*. The MFPT *τ*_12_(*θ*_1_) from starting position *θ*_1_ is then

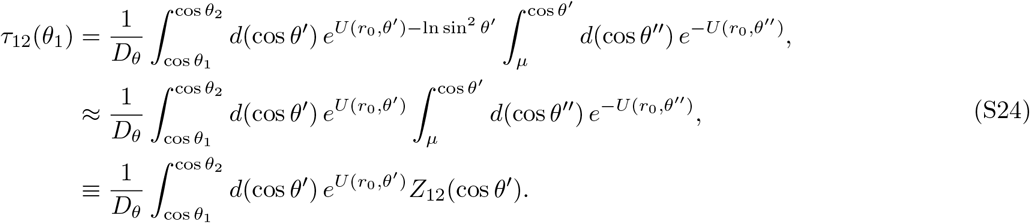

In the second line we have neglected the –ln sin^2^ *θ* contribution in the first exponential, since it does not significantly change the value of *τ*_12_(*θ*_1_), and in the third line we have introduced the function 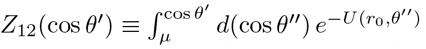. The final approximation is to note that *θ*_1_ is close to *θ*_max_, and *θ*_2_ is close to *θ*_min_, so we can replace cos *θ*_1_ in the integration bounds with *μ* = cos *θ*_max_, and replace cos *θ*_2_ with *v* = cos *θ*_min_. Thus the final integral for *τ*_12_(*θ*_1_) takes the form,

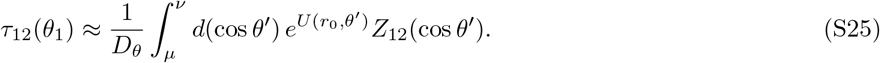

Since the angular dependence of *U*(*r_0_, θ’*) from Eqs. (S1)–(S2) is explicitly in terms of cos *θ’*, the integration variable, it turns out the integral in Eq. (S25) can be evaluated exactly, to yield a rather complex (but closed form) expression for *k*_12_,

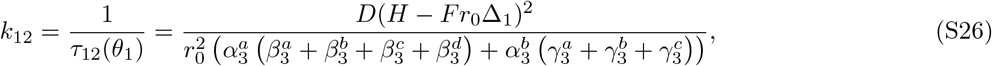

where,

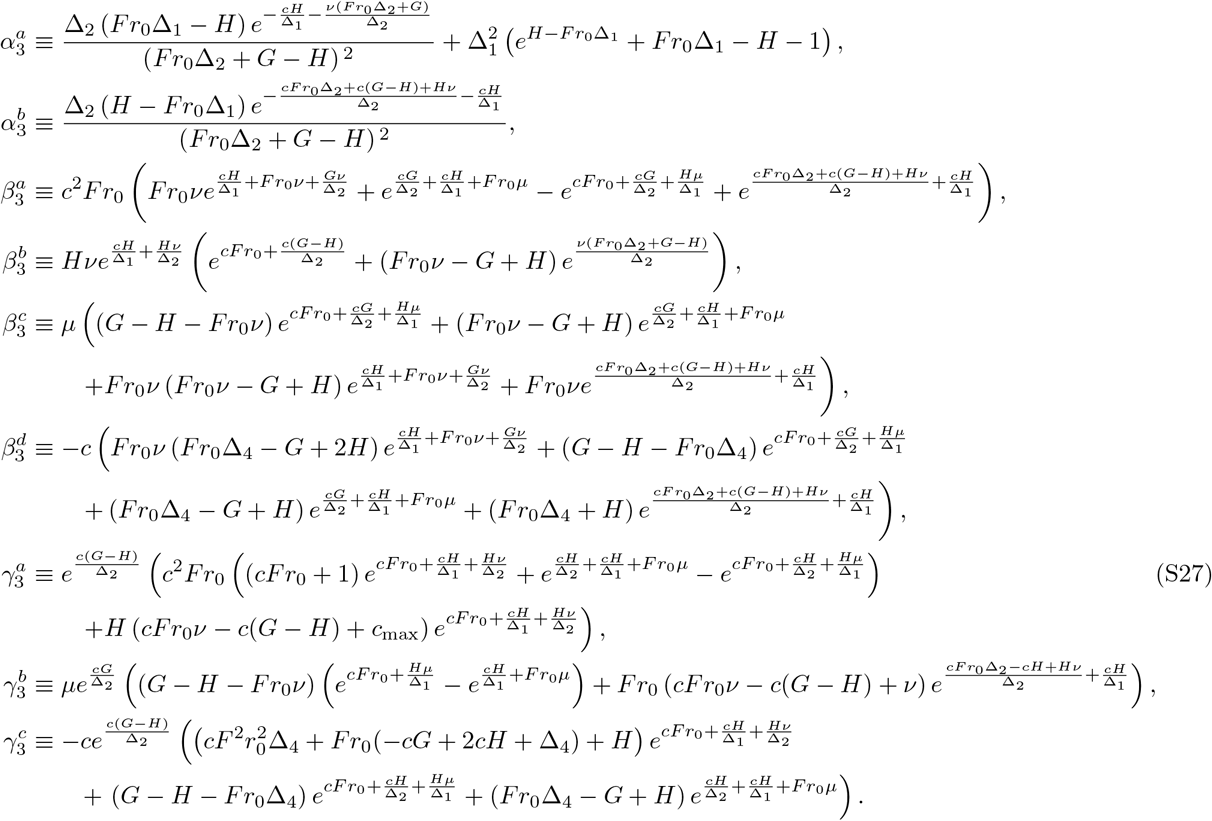

To get the transition rate *k*_21_ from state 2 back to 1, we note that a physically consistent model should relate *k*_12_ and *k*_21_ to each other through detailed balance. The quasi-equilibrium probability ratio of being in state 1 relative to state 2 (in the long-time limit, conditioned on the bond not rupturing), is approximately *Z*_12_(*c*)/*Z*_21_(*c*), where 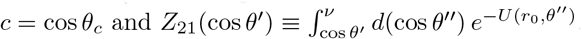. Thus we can write:

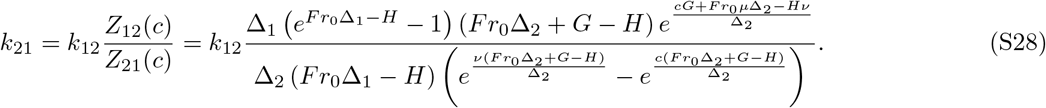

Eq. (S28), together with the expressions in Eqs. (S26)–(S27), gives a complete closed form result for *k*_21_.

### E. Survival probability and mean bond lifetime

The final part of the derivation involves expressing the survival probability Σ_*F*_ (*t*) and mean bond lifetime *τ*(*F*) in terms of the four rates *k*_10_, *k*_20_, *k*_12_, and *k*_21_, following a standard approach to first passage problems in discrete state kinetic networks [1]. As mentioned earlier, all these four rates are themselves functions of *F*, as can be seen in the results of the previous two sections, but for simplicity we do not show the *F* dependence explicitly.

Consider the probability *S_i_*(*t*) that the bond survived intact until time *t*, given that the system started in state *i* = 1, 2 at time *t* = 0. If we discretize time in infinitesimal steps of *δt*, with *t = nδt*, then the probability *S_1_(t)* can be written as

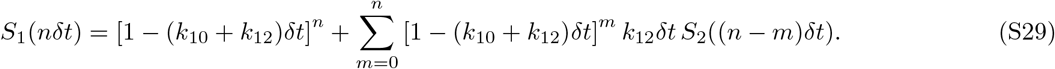

The right-hand side of Eq. (S29) can be understood as follows: *k*_10_*δt* is the probability to transition from 1 to 0 in time step *δt*, and *k*_12_*δt* is the probability to transition from 1 to 2 in time step *δt*. Thus we see that the first term on the right-hand side of Eq. (S29) is the probability that the bond survived without either rupturing or transitioning to state 2 for the entire *n* time steps. This is one contribution to *S_1_* (*nδt*). However there is another contribution, since the bond could still survive, but make at least one transition to state 2 during those *n* steps. The sum in Eq. (S29) is this second contribution, consisting of the cases where the bond does not leave state 1 for *m* time steps, then makes a transition to state 2, and survives the remaining *n* – *m* time steps. The last probability is just *S*_2_((*n – m*)*δt*). Taking the limit *δt* ⟶ 0, *n* = *t/δt* ⟶ ∞, we can rewrite Eq. (S29) as

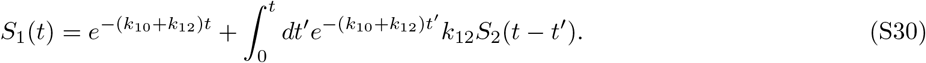

An exactly analogous argument for *S_2_(t)* yields a second integral equation,

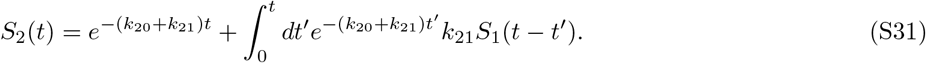

The system of equations, Eq. (S30)–(S31), can be solved by first applying a Laplace transform, 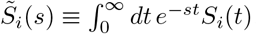. This gives

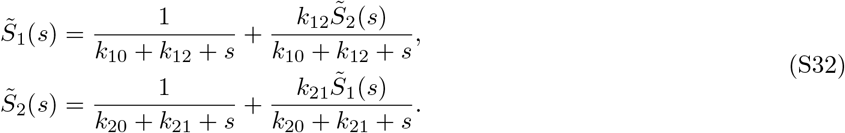

The solutions for 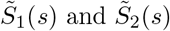 are then:

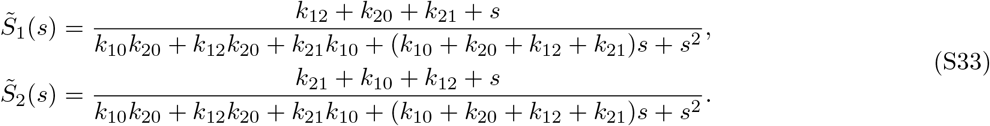

Before going further, note that if the system started in state *i* at time *t* = 0, the probability to rupture between times *t* and *t* + *δt* is just *S_i_(t) – S_j_(t + δt) ≈ – δt dS_i_(t)/dt*. Hence the mean time to rupture *τ_i_* given starting state *i* is just

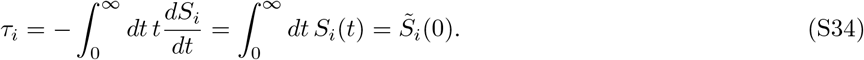

The second equality follows from integration by parts, and the fact that *S_i_*(0) = 1, *S_i_*(∞) = 0. Plugging *s* = 0 into Eq. (S33) thus gives

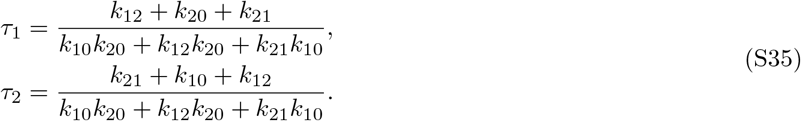

To get the final expression for the mean bond lifetime *τ*, we need the initial probabilities 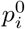 of being in state *i* at time *t* = 0. Since we assume the system has quasi-equilibrated at *F* = 0 before the application of force at *t* > 0, the ratio 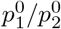 is just *Z*_12_(*c*)/*Z*_21_(*c*) evaluated at *F* = 0. From this we get the following probabilities:

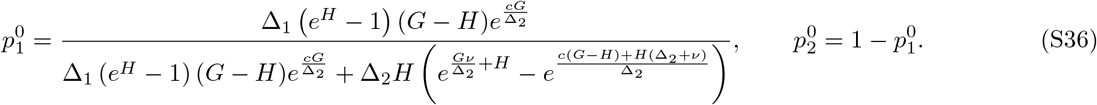

For the best-fit parameters given in Table 1 of the main text, these probabilities are 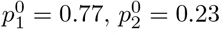. The initial small α probability was denoted as 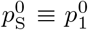 in the main text. The mean bond lifetime, weighting over all possible starting states, is given by

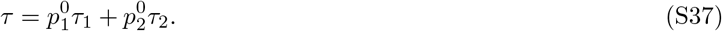

Eq. (S37), supplemented by Eqs. (S35)–(S36) and the expressions for *k*_10_, *k*_20_, *k*_12_, and *k*_21_ from the previous two sections (Eqs. (S15)–(S19), (S26)–(S28)), constitutes the complete theoretical result for *τ*.

Similarly the survival probability Σ*_F_(t)* is just the weighted sum of *S_1_(t)* and *S_2_(t)*,

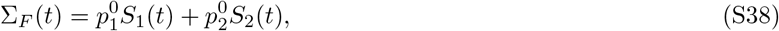

where *S_i_(t)* can be found by inverse Laplace transforming the solutions from Eq. (S33),

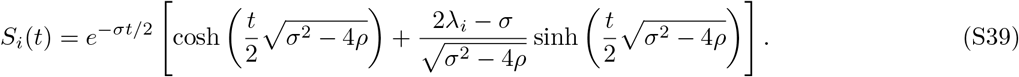

Here *ρ* ≡ *k*_10_*k*_20_ + *k*_12_*k*_20_ + *k*_10_*k*_21_, *σ* ≡ *k*_10_ + *k*_20_ + *k*_12_ + *k*_21_, λ_1_ ≡ σ – *k*_10_, λ_1_ ≡ *σ* – *k*_20_. Note that because the cosh and sinh share the same argument, Eq. (S39) can also be expressed in terms of two distinct exponential contributions with different prefactors. This is what leads to the double-exponential behavior seen in the survival probabilities in the main text.

Another aspect mentioned in the main text is the final conformational state of the system at the moment of rupture, whether it is state 1 (small *α*) or state 2 (large *α*). If the system could quasi-equilibrate at the applied force *F* before rupture occurred (i.e. if the rupture rates *k*_10_ and *k*_20_ were sufficiently small), the probability of being in state 1 at rupture would be *p*S = *k*_21_/(*k*_12_ + *k*_21_). For the case *F* = 15.1 pN, discussed in the main text, *p*S = 10^-4^ for the parameter values of Table 1. In reality, however, the system does not have time to fully quasi-equilibrate, and the actual probability of being in state 1 at rupture is 0.47 for this particular value of *F*. To derive this number, we define the splitting probability *π_i1_*, the probability that the system will rupture in state 1, given a starting state *i* at time *t* = 0. The splitting probabilities *π*_11_ and *π*_21_ satisfy the identities [1]:

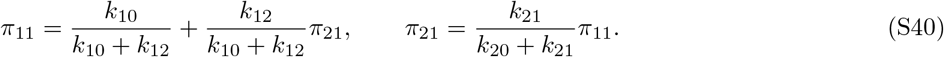

The first identity states the *π*_11_ involves two contributions: starting in state 1, the system can either rupture before jumping to state 2 (probability 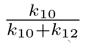) or jump to state 2 first (probability 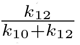) and then eventually make it back to state 1 to rupture (probability *π*_21_). Similarly for the second identity, *π*_21_ is equal to the probability of jumping to state 1 before rupture 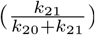 times *π*_11_. Eq. (S40) can be solved for *π*_11_ and *π*_21_,

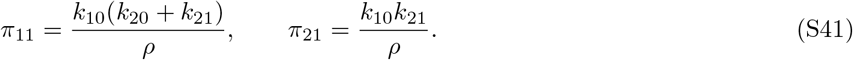

The probability of rupturing from state 1 independent of initial state is 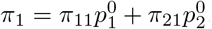. For *F* = 15.1 pN we get *π*_1_ = 0.47, which was denoted as *π*_S_ ≡ *π*_1_ in the main text.

The final quantity discussed in the main text is *τ_L_*, the mean duration of the large α conformation (state 2), measured from the first entrance into the state until either rupture occurs or the system transitions to state 1. Since the total escape rate from state 2 is *k*_20_ + *k*_21_, the probability of leaving state 2 between times *t* and *t + δt*, where *t* = 0 is the time of entrance, is: *δt*(*k*_20_ + *k*_21_) exp(–(*k*_20_ + *k*_21_)*t*). Thus the mean duration is:

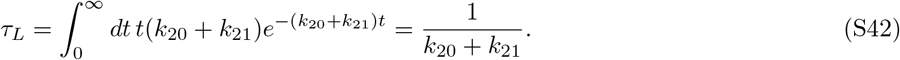

## II. CONSISTENCY WITH AN EARLIER CATCH BOND MODEL

One of the nice features of our approach is its generality: even though we focus in the main text on a system with a substantial angular barrier *H*, the theoretical model continues to hold even in the absence of such a barrier. In this section we show that the *τ*(*F*) expression for an earlier, barrier-less model for catch bonds in selectin systems, introduced and numerically verified in Ref. [2], is just a special case of our more general *τ*(*F*).

For the selectin case, the full angular range was used, so *θ*_min_ = 0° and *θ*_max_ = *π*. There was no angular barrier or energy offset, so *H = G* = 0, and we can take *θ_c_* = 90°, since the border between state 1 and state 2 is arbitrary without a barrier present. In this case from Eq. (S17) we see that 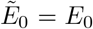 and 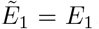. Plugging all these values into the expressions for the transition rates derived above, we find relatively simple results:

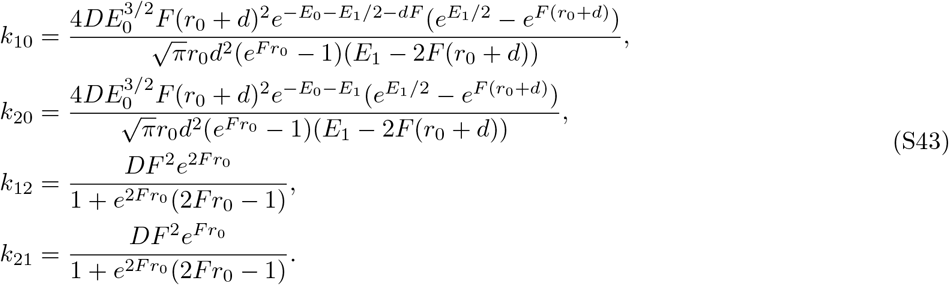

Without an angular barrier the transitions between angular regions are many orders of magnitude faster than the transitions to rupture, as can be seen from the fact that *k*_10_ and *k*_20_ both include a factor of *e^-E_0_^* in the numerator that is not present in *k*_12_ and *k*_21_. Typically the factor *e^-E_0_^* ≪ 1 since *E*_0_ sets the overall energy scale for rupture, and *E*_0_ ~ 17 – 26 (units of *k_B_T*) for the systems considered in Ref. [2]. Hence we can assume in this case that *k*_10_, *k*_20_ ≪ *k*_12_, *k*_21_. This simplifies the expressions for *τ*_1_ and *τ*_0_ in Eq. (S35), so that *τ*_1_ ≈ *τ*_0_ ≈ (*k*_12_+*k*_21_)/(*k*_12_*k*_20_+*k*_21_*k*_10_). Hence the mean bond lifetime is also the same as *τ*_1_ and *τ*_2_,

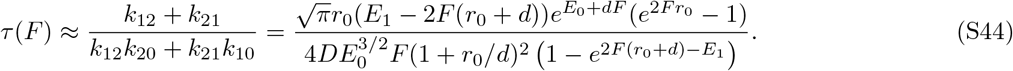

This is in complete agreement with the *τ*(*F*) from Eq. (2) in Ref. [2]. The survival probability in this limit becomes a single exponential, Σ_*F*_(*t*) ≈ exp(–*t/τ(F)*), with *τ*(*F*) from Eq. (S44).

Our derivation of *τ*(*F*) and Σ_*F*_ (*t*) in the previous section also allows us to make a comparison to another catch bond model. By partitioning the parameter space into two angular states, and focusing on four transition rates (*k*_10_, *k*_20_, *k*_12_, *k*_21_), our approach on the surface seems analogous to the phenomenological two-state catch bond model [3, 4]. However in this phenomenological model each transition rate *k_ij_* is assumed to have a simple Bell-like dependence on the force, 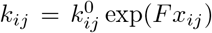, for coefficients 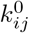 and distances *x_ij_*. Our general expressions for the transition rates in the previous sections, and even the simplified versions of Eq. (S43) in the barrier-less limit, are quite different from Bell models. This is because these rates are derived from an underlying energy landscape based on a structural model. Our parameters thus directly connect to structural / energetic features of the system, in contrast to the 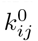, *x_ij_* parameters of the phenomenological model.

## III. MAXIMUM LIKELIHOOD FITTING TO THE EXPERIMENTAL DATA

The experimental data 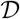 (Ref. [5] Fig. 4A) consists of *N* = 803 points, 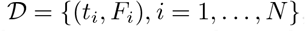, where *t_i_* is the measured bond lifetime, and *F_i_* is the applied force. Let ⋀ be the set of free parameters in the model other than *F*. The probability 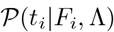 of observing the *i*th bond lifetime, given force *F*_i_ and particular set of parameter values ⋀, is:

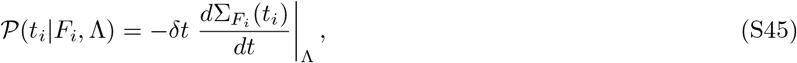

where Σ*_F_(t)* is the survival probability at force *F*. Since we have an analytical expression for Σ*_F_(t)* from Eqs. (S38)–(S39), we also can get an analytical form for *dΣ_F_(t)/dt*, which allows us to evaluate 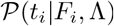. The joint probability of the entire data set, given the model parameters, is

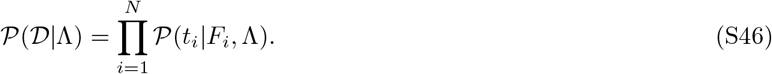

To find the best-fit parameter set ⋀, we maximize the log-likelihood function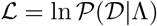,

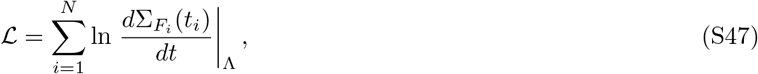

where we have neglected an additive constant dependent on *δt* that does not affect the fitting.

To prevent the maximization algorithm, implemented in *Mathematica*, from veering into unphysical regions of parameter space, the parameters were constrained to vary over physically sensible ranges: *E*_0_, *H, G, d, r*_0_ ≥ 0, *α*_max_ > *α_c_* > *α*_min_ + *γ*. Here the buffer angle *γ* was set to 5°, to put a constraint on the minimum possible angular range for the small α conformational state. This choice of *γ* was based on the magnitude of fluctuations in molecular dynamics trajectories of *α* in Ref. [6], though other choices of γ within a few degrees also lead to similar maximum log-likelihoods and best-fit parameter sets ⋀.

